# Hemoglobin alpha is a redox-sensitive mitochondrial-related protein in T-lymphocytes

**DOI:** 10.1101/2024.09.16.613298

**Authors:** Emily C. Reed, Valeria A. Silva, Kristen R. Giebel, Tamara Natour, Tatlock H. Lauten, Caroline N. Jojo, Abigail E. Schleiker, Adam J. Case

## Abstract

Hemoglobin subunits, which form the well-characterized, tetrameric, oxygen-carrying protein, have recently been described to be expressed in various non-canonical cell types. However, the exact function of hemoglobin subunits within these cells remains to be fully elucidated. Herein, we report for the first time, the expression of hemoglobin alpha-a1 (Hba-a1) in T-lymphocytes and describe its role as a mitochondrial- associated antioxidant. Within naïve T-lymphocytes, Hba-a1 mRNA and HBA protein are present and highly induced by redox perturbations, particularly those arising from the mitochondria. Additionally, preliminary data using a T-lymphocyte specific Hba-a1 knock-out mouse model indicated that the loss of Hba-a1 led to an exacerbated production of mitochondrial reactive oxygen species and inflammatory cytokines after a stress challenge, further supporting the role of HBA acting to buffer the mitochondrial redox environment. Interestingly, we observed Hba-a1 expression to be significantly upregulated or downregulated depending on T-lymphocyte polarization and metabolic state, which appeared to be controlled by both transcriptional regulation and chromatin remodeling. Altogether, these data suggest Hba-a1 may function as a crucial mitochondrial-associated antioxidant and appears to possess critical and complex functions related to T- lymphocyte activation and differentiation.

## Introduction

The extensive, genetically-conserved history of hemoglobin and its function as an oxygen carrier has been well known since the early 19^th^ century [1]. This protein was classically believed to be exclusive to erythrocytes until the late 20^th^ century, when it was first discovered to be expressed in macrophages [2]. Since then, hemoglobin has been discovered to be expressed in many other non-canonical cell types including mesangial cells [3], hepatocytes [4], alveolar epithelium [5–7], vascular endothelium [8], neurons [9–11], chondrocytes [12], retinal cells [13, 14], cervical cells [15], endometrial cells [16], and cardiac cells [17]. This ever-growing list suggests the presence of intracellular hemoglobin within a specific cell type is no longer an atypical observation. For seemingly a ubiquitous protein, the function of hemoglobin within this expansive range of cell types is quite diverse. In the few accounts that have attempted to elucidate the function of hemoglobin in non-erythrocyte cells, its functions have been reported as a pseudo-peroxidase [18, 19], nitric oxide regulator [8, 20], glycation mediator [21], as well as responsive to hypoxia [6, 12]. While varied, a common theme emerges among these functions: redox modulation.

While it is becoming accepted that reactive oxygen species (ROS) and the redox environment are crucial to cellular signaling as opposed to simply a detrimental ‘side-effect’ of metabolism and stress, very little is known about redox signaling in adaptive immunity, particularly T-lymphocytes. It was only in the early 2000’s where it was discovered that ROS, produced by a phagocytic-like NADPH oxidase, are critical signaling molecules for T-lymphocyte activation upon receptor stimulation [22, 23]. Moreover, the importance of mitochondrial ROS in T-lymphocyte development and function only began in the early 2010’s [24–28]. Currently, it is clear that these various sources of ROS are indeed essential for T-lymphocyte activation and effector functions, as well as they appear to be closely aligned with the metabolic state of these cells [29]. As T-lymphocytes activate and differentiate, their metabolic utilization dramatically shifts between glycolysis and oxidative phosphorylation dependent upon polarization stage, which simultaneously alters the cellular redox environment [30, 31].

However, the understanding of how the redox and metabolic environments are controlled in these different stages of T-lymphocyte activation remains unknown.

Herein, we report the first observation of hemoglobin alpha expression in both murine and human T- lymphocytes. Similar to other studies of non-canonical hemoglobin expression, T-lymphocyte hemoglobin appears highly responsive to pro-oxidant stimuli, particularly mitochondrial ROS. Moreover, the loss of hemoglobin alpha in T-lymphocytes elevates the levels of mitochondrial ROS and decreases the proportion of naïve cells in the populations, suggesting accelerated activation. Additionally, the regulation of hemoglobin in these adaptive immune cells is highly dynamic and complex, demonstrating robust downregulation and upregulation that appears both dependent upon the metabolic/redox state of the mitochondria as well as under various expression control mechanisms (i.e., transcriptional regulation and chromatin remodeling). Overall, this work has identified hemoglobin alpha as a redox-responsive protein that is closely aligned to mitochondrial function in T-lymphocytes, which has important implications regarding our fundamental understanding of how the redox environment shapes T-lymphocyte development, activation, and differentiation.

## Materials and Methods

### Mice

Wild-type C57BL/6J (#000664; shorthand WT) and CD4 cre (#022071; shorthand CD4-cre) mice were obtained from Jackson Laboratories (Bar Harbor, ME, USA). Estrogen receptor 1 alpha cre (Esr1-cre) mice on a CD1 background were generated and bred in house as previously described [32]. Conditional manganese superoxide dismutase (MnSOD) knock-out mice were bred in house as previously described [24, 26].

Conditional mitochondrial catalase expression mice were graciously provided by Dr. Holly Van Remmen as previously described [33]. Whole body and conditional Hba-a1 knock-out mice were graciously provided by Dr. Brant Isakson as previously described [20]. All conditional knock-out or over-expression mice were crossed with CD4-Cre mice to generate T-lymphocyte-specific modified progeny. All mice were bred in house to eliminate shipping stress and microbiome shifts, as well as co-housed with their littermates (≤5 mice per cage) prior to the start of experimentation to eliminate social isolation stress. Mice were housed with standard pine chip bedding, paper nesting material, and given access to standard chow (#8604 Teklad rodent diet, Inotiv, West Lafayette, IN, USA) and water ad libitum. Male and female experimental mice between the ages of 8-12 weeks were utilized in all experiments, but no sex differences were observed so data are presented as pooled independent of sex. Experimental mice were randomized, and when possible, experimenters were blinded to the respective cohorts until the completion of the study. Mice were sacrificed by pentobarbital overdose (150 mg/kg, Fatal Plus, Vortech Pharmaceuticals, Dearborn, MI, USA) administered intraperitoneally. All mice were sacrificed between 7:00 and 9:00 Central Time to eliminate circadian rhythm effects on T-lymphocyte function. All procedures were reviewed and approved by Texas A&M University Institutional Animal Care and Use Committees.

### Repeated social defeat stress paradigm

Repeated social defeat stress (RSDS) was performed as described in [32]. Briefly, chemogenetically-altered Esr1-cre mice were injected intraperitoneally with clozapine-N-oxide (1 mg/kg) to induce aggressive behaviors towards both male and female experimental mice. For RSDS, the experimental mouse was placed into the aggressor’s cage for 1 minute, during which the aggressor mouse physically confronts and induces a traumatic fear response in the experimental mouse. Following this interaction, a clear, perforated barrier was placed in between the two mice and the mice were co-housed for 24 hours. This process was repeated for 5 consecutive days. Control mice were conspecifically pair-housed for the duration of the protocol with no exposure to aggressive mice. Experimental mice were monitored for any signings of wounding and lameness, and were removed from the study if exclusion criteria were met (wounds >1 cm, presence of any lameness).

### In-vivo LPS administration

Lipopolysaccharide from Salmonella Minnesota (#89152-786, VWR) was diluted with sterile 1X PBS and administered in one dose intraperitoneally at 5 mg/kg and sacrificed 6 hours later. Dose was chosen based on previous work [34, 35].

### In-vivo NAC administration

1% N-acetyl cysteine (NAC) was supplemented in the drinking water of mice starting 1 day before RSDS, and was supplied for the duration of the experiment (7 days total). NAC was dissolved in 4% sucrose water to mask taste, and control mice received 4% sucrose water only. Fresh water was made every three days and bottles were weighed daily to ensure equal consumption between control and NAC groups.

### Mouse T-lymphocyte isolation, culture, and activation

Splenic T-lymphocytes were isolated using negative magnetic selection as previously described [27]. Briefly, spleens were collected and disrupted into a single cell suspension and passed through a 70 μM nylon mesh filter (#22363548, ThermoFisher Scientific). Red blood cells were removed using red blood cell lysis buffer (150 mM NH_4_Cl, 10 mM KHCO_3_, 0.1 mM EDTA). T-lymphocytes were negatively selected using the EasySep Mouse total, CD4+, CD8+, or CD4+ memory T-cell isolation kit (StemCell Technologies #19851, #19852, #19853, #19767) per manufacturer’s instructions. T-lymphocytes were counted, and viability was assessed using Trypan Blue exclusion on a Bio-Rad TC20 Automated Cell Counter. For activation, cells were plated at 800,000 cells/mL with anti-CD3/28 Dynabeads (Dynabeads, #11456D) in a 1:1 cell to bead ratio in T-lymphocyte media consisting of RPMI media supplemented with 10% Fetal Bovine Serum, 2 mM Glutamax, 10 mM HEPES, 100 U/mL penicillin/streptomycin and 50 μM of 2-mercaptoethanol. Cells were cultured in 5% CO_2_, 37**°**C incubator (HERAcell Vios 160i CO_2_ incubator, ThermoFisher Scientific). Where indicated, cells were activated with 10ng/mL phorbol myristate acetate (PMA) and 500ng/mL ionomycin for 24 hours at 5% CO_2_, 37**°**C incubator.

### Human T-lymphocytes

Human peripheral blood pan T-lymphocytes (#70024.1, StemCell Technologies) were thawed and cultured in the aforementioned T-lymphocyte media. Naïve T-lymphocytes were cultured for 24 hours and treated with 50- 200uM of H_2_O_2,_ and harvested for subsequent protein or RNA extraction as outlined below.

### Cell lines

Mouse T-lymphoblast cell line TK1 (CRL-2396, ATCC) was cultured in the aforementioned T-lymphocyte media. TK1 cells were cultured in a 5% CO_2_, 37**°**C incubator, and sub-cultured according to ATCC instructions to avoid over-confluence.

### Hba-a1 lentivirus over-expression transfection

Recombinant lentivirus containing Hba-a1 exons (pLV[Exp]-hPGK>mHba-a1[NM_008218.2]-EF1A>EGFP (Vector ID: VB240229-1499mdx)) and control GFP virus (pLV[Exp]-EF1A>EGFP (Vector ID: VB900088- 2243bzq) was created by VectorBuilder. Cells were transfected by the addition of virus at various MOIs and cultured 24 hours at 5% CO_2_, 37**°**C incubator.

### T-lymphocyte polarization

CD4+ T-lymphocytes were isolated and polarized to T_H_1, T_H_2, T_H_17, T_reg_ cells by activation with Dynabeads anti-CD3/28 beads in a 1:1 cell to bead ratio and cytokine supplementation. To polarize to various subtypes, CD4+ T-lymphocytes were supplemented with the following: T_H_1: 15 ng/mL IL-12 (StemCell Technologies #78028.1), 150 ng/µLIL-2 (StemCell Technologies #78081), 5 ug/mL anti-IL-4 (clone 11B11, Miltenyi #130-095-709); T_H_2: 10 ng/mL IL-4 (StemCell Technologies #78047.1), 150 ng/µLIL-2, 5 ug/mL anti-IFNγ (clone XMG1.2, Miltenyi #130-095-729); T_reg:_ 150 ng/µL IL-2, 15 ng/mL TGFβ (Miltenyi #130-107-758), 10 ug/mL anti-IL-4, 10 ug/mL anti-IFNγ. CD4+ T-lymphocytes were polarized to T_H_17 cells using the CytoBox T_H_17 mouse kit (Miltenyi #130-107-758) and supplemented according to manufacturer’s instructions. Cells were cultured for 72 hours before analysis. CD8+ T-lymphocytes were isolated and polarized to T_mem_ by culturing with 50ng/mL IL-2 and 1:1 cell to Dynabead ratio for 3 days, harvested and replated at 100,000 cells/well and treated with 50ng/mL IL- 2 or 10ng/mL IL-15. Cells were harvested 4 days later and immunophenotyped via flow cytometry.

### In vitro treatments

Hydrogen peroxide, Auranofin (#102988-762, Avantor), 5,6-Dichlorobenzimidazole 1-β-D-ribofuranoside (DRB, D1916, Sigma Aldrich), and actinomycin D (#A9415, Sigma Aldrich) were treated at doses indicated.

Concentrations of all drugs were chosen on prior dose curves or previous work [36, 37]. Hypoxia induction was performed by placing cells in a hypoxia incubator chamber (#27310, StemCell) with 1% O_2_ at 37**°**C incubator for 6 hours.

### Flow cytometry immunophenotyping and redox assessment

T-lymphocytes were immunophenotyped via 4-laser Attune NxT flow cytometer (ThermoFisher Scientific) as previously described [27]. Cells were stained with 1:1000 dilutions of CD3ε PE-Cy7 (#25-0031-82, ThermoFisher Scientific), CD4 eFluor 506 (#69004182, ThermoFisher Scientific), and CD8 Super Bright 702 (#67008182, ThermoFisher Scientific) antibodies along with 1 μM MitoSOX Red (#M36008, ThermoFisher Scientific) in RPMI media to assess mitochondrial reactive oxygen species in various T-lymphocyte subpopulations. Mean fluorescence intensity (MFI) of MitoSOX Red was reported as a readout of mitochondrial ROS levels. 100nM Tetramethylrhodamine ethyl ester (TMRE) (#T669, ThermoFisher Scientific) MFI was used as a measure of actively metabolizing mitochondria.

### RNA extraction, cDNA production, and quantitative real-time RT-PCR

T-lymphocyte RNA isolation and gene expression was assessed as previously described [38]. Briefly, mRNA was extracted using a RNAeasy plus mini kit (#74136, Qiagen) and quantified using NanoDrop One Spectrophotometer (#13400518, Thermo Scientific). RNA was then transformed into cDNA using ThermoFisher High-Capacity cDNA Reverse Transcriptase Kit (#4374967, Applied BioSystem). Generated cDNA was used for real time quantitative PCR. Primers for genes of interest were designed using NIH primer-BLAST spanning exon-exon junctions. Cq values were determined, and relative gene expression was calculated by comparing housekeeping 18s ribosomal gene expression to gene of interest (2^-ddCq).

### Protein analysis

Protein was isolated from T-lymphocytes using RIPA lysis buffer (#PI89900, Thermo Scientific) and 1% Halt protease inhibitor cocktail (#PI87785, Thermo Scientific). Samples were subsequently subjected to sonication and centrifugation to obtain soluble protein, which was quantified using Pierce BCA protein assay kit (#PI23227, Thermo Scientific). Mouse and human hemoglobin alpha protein was assessed via Total Protein Jess Automated Western Blot (Bio-techne) as described in [38] using anti-HBA primary antibody - Rabbit (#PIPA579347, Fisher) 1:20. Analysis was performed using Compass Software for Simple Western. Protein was also assessed by tandem mass spectrometry for the presence/absence of hemoglobin alpha using the University of Nebraska Medical Center Mass Spectrometry and Proteomics Core facility.

### Chromatin Accessibility

Chromatin was isolated from primary and activated mouse T-lymphocytes with EpiQuik Chromatin Accessibility Assay (#P-1047-48, Epigentek) according to manufacturer’s instructions. Chromatin regions of interest (i.e., Hba-a1 promoter) were then assessed for accessibility via RTqPCR.

### Seahorse Mitochondrial Stress Test

T-lymphocyte mitochondrial metabolism assessment was performed as previously described in [38]. Briefly, TK1 cells were plated in Seahorse XF RPMI media (#103576-100, Agilent) supplemented with 10mM Seahorse XF Glucose (#103577-100, Agilent), 1mM Seahorse XF Pyruvate (#103578-100, Agilent), 2mM Seahorse XF L-Glutamine (#103579-100, Agilent). Cells were adhered to seahorse cell microplates using 1ug/cm^2 Cell-Tak (#354240, Corning) and seeded at a density of 250,000 cells per well. Mitochondrial inhibitors (1 µM Oligomycin, 1µM 4-trifluoromethoxy-phenylhydrazone (FCCP), 0.5µM Rotenone and Antimycin, # 103015-100, Agilent), were injected into each well and oxygen consumption rate (OCR) was measured via Seahorse XFe96 Analyzer.

### Statistical analysis

All data presented as mean ± standard error of the mean (SEM), with N values listed in figure legends. Normality was assessed using Shapiro-Wilk normality test before statistical analysis. For two group comparisons, Mann-Whitney U test or Student’s t-test were utilized. For experiments with 3 or more groups, an ordinary one-way ANOVA was performed. Experiments containing two categorical groups were assessed using two-way ANOVA. All statistics were completed using GraphPad Prism version 10.1.2.

## Results

### Hemoglobin alpha (Hba-a1) is expressed and functional in T-lymphocytes

To our knowledge, the only previous report of hemoglobin alpha in T-lymphocytes was from one of our own experiments where mice with elevated levels of T-lymphocyte mitochondrial ROS, due to the conditional loss of manganese superoxide dismutase (MnSOD), showed elevated levels of Hba-a1 mRNA by microarray analysis [24]. Since that analysis was performed on bulk pan-T-lymphocytes from MnSOD knock-out mice, there was the possibility that this observation of hemoglobin alpha was simply due to erythrocyte contamination.

However, we have recently performed a single cell RNA sequencing analysis on T-lymphocytes from a mouse model of psychological trauma (repeated social defeat stress; RSDS), which we have shown elevates mitochondrial ROS in these cells, and again, demonstrated robust and significant elevation of Hba-a1 mRNA in both CD4+ (+23 fold, p=7.09e^-17^) and CD8+ T-lymphocytes (+25 fold, p=4.21e^-11^) [38, 39]. We have confirmed these original screening observations of Hba-a1 mRNA levels in both MnSOD knock-out T-lymphocytes and RSDS T-lymphocytes via RT-qPCR (**Figure 1A-B**). To extend these findings and investigate whether Hba-a1 upregulation in T-lymphocytes could be elicited by an immunological challenge, LPS was administered to mice. T-lymphocytes from LPS-treated mice again displayed significant upregulation of Hba-a1 mRNA compared to heathy controls (**Figure 1C)**. To confirm the presence of HBA protein, T-lymphocyte protein was assessed by mass spectrometry, and not only confirmed the presence of HBA protein in T-lymphocytes but also its significant increase after RSDS (**Figure 1D**). Together, these data indicate that T-lymphocytes express hemoglobin alpha at both the RNA and protein level, as well as demonstrate its dynamic regulation to various stimuli. Next, to preliminarily assess if hemoglobin alpha possessed functionality in T-lymphocytes, we performed RSDS on mice lacking Hba-a1 constitutively as well as specifically in T-lymphocytes. Strikingly, Hba-a1 loss aggravated the RSDS-induced level of mitochondrial ROS (**Figure 1E-F**), suggesting Hba-a1 may possess mitochondrial antioxidant properties. Moreover, previous work from our lab and others has demonstrated that mitochondrial ROS are essential for T-lymphocyte activation [24–28], and indeed, Hba-a1 loss concurrently led to decreased numbers of naïve T-lymphocytes and elevated levels of pro-inflammatory interleukin 6 (IL-6) in these mice (**Figure 1G-H**), suggesting altered activation in these cells. These pilot data establish a functional role for hemoglobin alpha in buffering the redox environment and activation of T-lymphocytes, which warrants further investigation.

**Figure 1.**
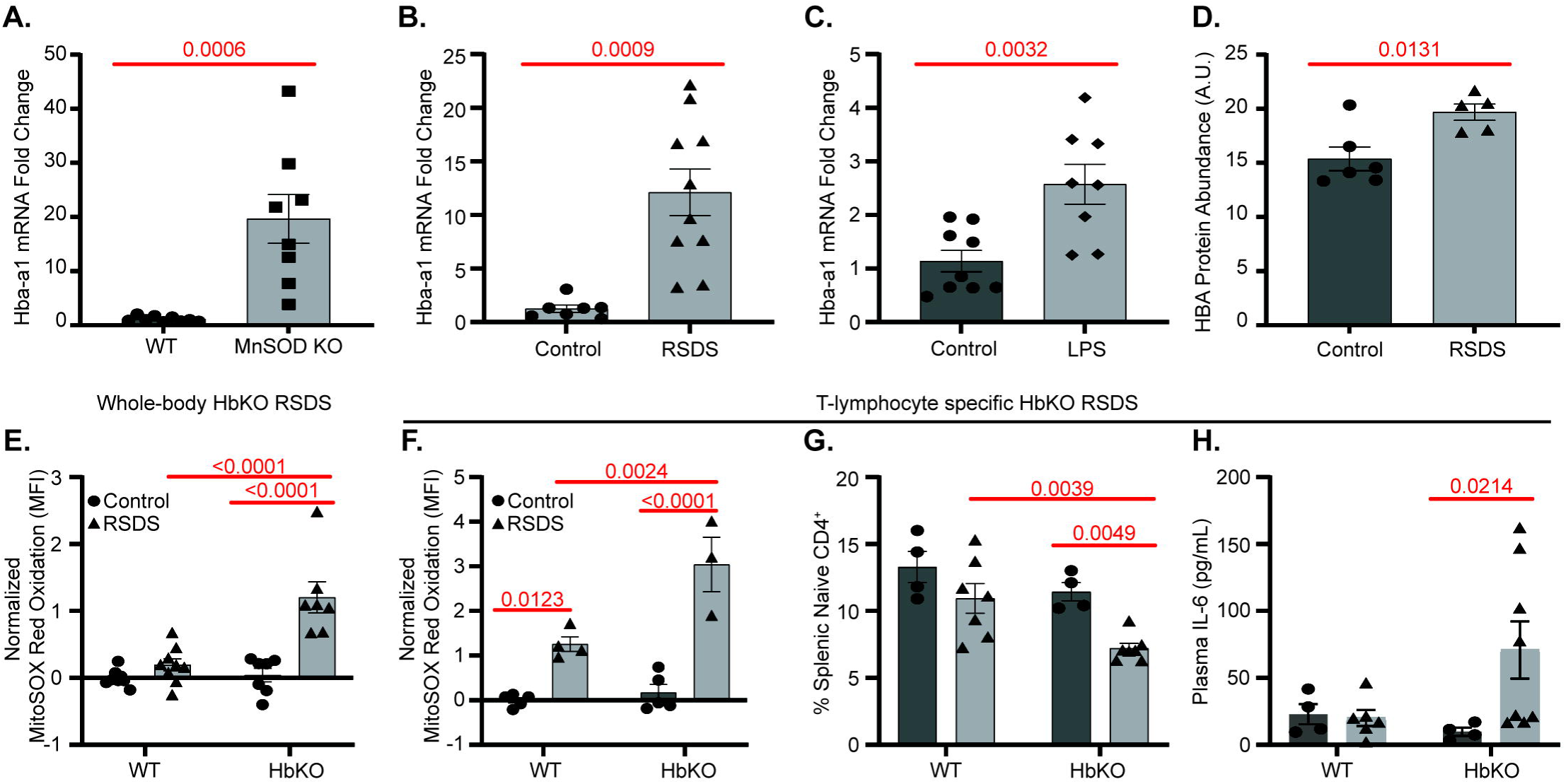
Hba-a1 is present in and impacts T-lymphocyte function. **A**. Hba-a1 mRNA assessed by RTqPCR in splenic T-lymphocytes from wild-type mice (WT) or mice lacking MnSOD (MnSOD KO) specifically in T-lymphocytes. **B**. Hba-a1 mRNA assessed by RTqPCR in splenic T-lymphocytes from mice exposed to repeated social defeat stress (RSDS). **C**. Hba-a1 mRNA assessed by RTqPCR in splenic T-lymphocytes from mice treated with 5 mg/kg LPS. **D**. HBA protein assessed by mass spectrometry in splenic T-lymphocytes from mice exposed to repeated social defeat stress (RSDS). **E**. MitoSOX Red fluorescence assessed by flow cytometry in splenic T-lymphocytes from WT and whole-body Hba-a1 knock-out (HbKO) mice after RSDS. Data normalized relative to WT control animals. **F**. MitoSOX Red fluorescence assessed by flow cytometry in splenic T-lymphocytes from WT and T-lymphocyte-specific HbKO mice after RSDS. Data normalized relative to WT control animals. **G**. Percentage of naïve (CD62L+, CD44-) CD4+ splenic T-lymphocytes assessed by flow cytometry from WT and T-lymphocyte-specific HbKO mice after RSDS. **H**. Plasma interleukin 6 (IL-6) levels assessed by U-Plex Mesoscale from WT and T-lymphocyte-specific HbKO mice after RSDS. Statistics by Student’s t-test (**A-D**) or 2-way ANOVA with Tukey’s post-hoc analysis (**E-H**).

### Hba-a1 is redox-responsive in T-lymphocytes

Data from our lab has consistently shown an increase of mitochondrial ROS in T-lymphocytes after RSDS [38–41]. Given that Hba-a1 is also upregulated after RSDS, we investigated if ROS perturbations could directly induce Hba-a1 expression in T-lymphocytes. Primary murine T-lymphocytes were treated with hydrogen peroxide (H_2_O_2_), and indeed demonstrated significant induction of both Hba-a1 mRNA and HBA protein post- H_2_O_2_ treatment (**Figure 2A-B**). This exact phenomenon was repeated in human T-lymphocytes (**Figure 2C-D**), showing that human T-lymphocytes also express redox-sensitive hemoglobin alpha mRNA and protein. Next, T-lymphocyte cell lines were screened for the presence of hemoglobin alpha. Interestingly, we only identified Hba-a1 mRNA expression in the mouse T-lymphocyte cell line TK1 (**Figure 2E**), whereas in EL4 (mouse T- lymphocytes) and Jurkat (human T-lymphocytes) cell lines, the levels of hemoglobin alpha mRNA expression were undetectable (data not shown). Furthermore, Hba-a1 mRNA in TK1 cells showed redox-sensitivity to H_2_O_2_ similar to primary mouse and human T-lymphocytes (**Figure 2E**). Using both primary mouse T- lymphocytes and TK1 cells to further explore Hba-a1 expression in T-lymphocytes, we observed an increase in Hba-a1 mRNA with Auranofin (AFN; a thioredoxin reductase inhibitor) treatment, demonstrating perturbations of endogenous H_2_O_2_ degradation could also modulate Hba-a1 expression (**Figure 2F-G**). Furthermore, hypoxia (which is known to elevate mitochondrial ROS [42, 43]) also was sufficient in increasing Hba-a1 mRNA in both primary mouse T-lymphocytes and TK1 cells (**Figure 2H-I**). Conversely, known hemoglobin/heme inducers (i.e., hemin and erythropoietin (EPO)) and catecholamine neurotransmitters (i.e., dopamine, norepinephrine, and epinephrine; relevant to psychological trauma [32, 39, 41]) failed to induce T-lymphocyte Hba-a1 mRNA expression (**Supplemental Figure 1A-C**), suggesting specific redox-regulatory control of hemoglobin alpha in T-lymphocytes. To further support this notion of redox-regulation, antioxidant administration (drinking water supplemented with N-acetyl cysteine) and genetic over-expression of mitochondrial-targeted catalase significantly attenuated RSDS-induced Hba-a1 elevations in T-lymphocytes (**Figure 2J-K**). Together, these data put forth strong evidence that hemoglobin alpha is redox-regulated, particularly by H_2_O_2_, in T-lymphocytes.

**Figure 2.**
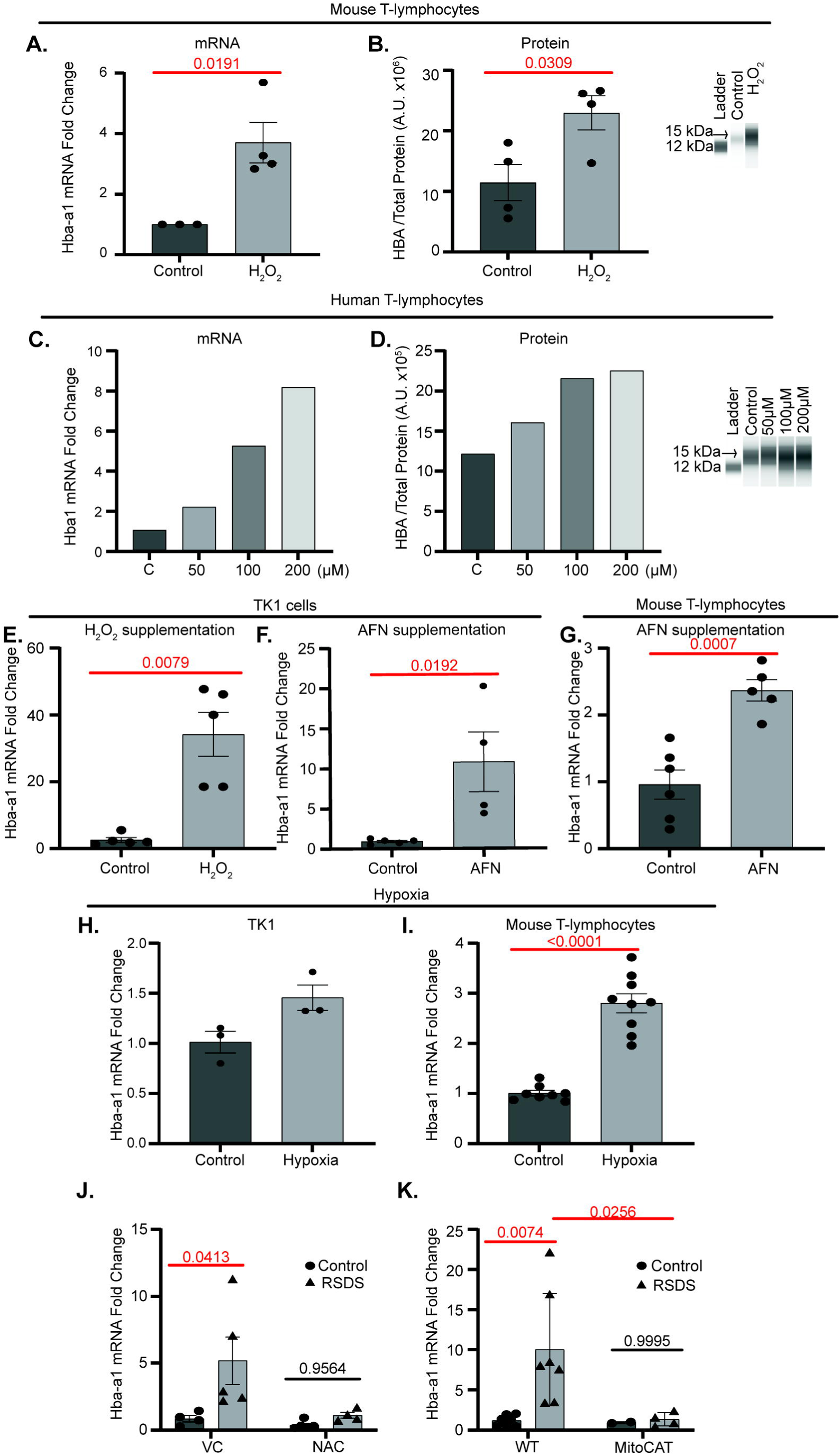
T-lymphocyte hemoglobin alpha is redox responsive. A-B. Mouse naïve splenic T-lymphocytes treated with 50µM of hydrogen peroxide (H_2_O_2_) for 24 hours. Hba-a1 mRNA assessed by RTqPCR and HBA protein assessed by Jess Automated Western. **C-D**. Human naïve T-lymphocytes treated with various concentrations of H_2_O_2_ for 24 hours. Hba-a1 mRNA assessed by RTqPCR and HBA protein assessed by Jess Automated Western. **E**. TK1 cells treated with 250µM H_2_O_2_ for 6 hours and Hba-a1 mRNA assessed by RTqPCR. **F**. TK1 cells treated with 1 µM Auranofin (AFN) for 12 hours and Hba-a1 mRNA assessed by RTqPCR. **G**. Mouse naïve splenic T-lymphocytes treated with 1 µM AFN for 24 hours, and Hba-a1 mRNA assessed by RTqPCR. TK1 cells (**H**.) and primary T-lymphocytes (**I**.) subjected to 1% O_2_ for 6 hours assessed for Hba-a1 mRNA by RTqPCR. **J**. Mice given water supplemented with sucrose (VC: vehicle control) or N- Acetyl Cysteine (NAC) and Hba-a1 mRNA assessed after RSDS by RTqPCR. **K**. Wild-type (WT) and T- lymphocyte specific mitochondrial-targeted catalase mice (MitoCAT) Hba-a1 mRNA assessed after RSDS. Statistics by Student’s t-test (**A-I**) or 2-way ANOVA with Tukey’s post-hoc analysis (**J-K**).

### Hba-a1 expression is altered in various T-lymphocyte subtypes

The previous experiments were mainly performed on primary splenic T-lymphocytes isolated from mice, which are predominantly in a naïve state, so we next queried how Hba-a1 expression may be altered in different states of activation and polarization. First, we activated primary mouse T-lymphocytes using anti-CD3/28 crosslinking antibodies, and surprisingly, observed a rapid decrease in Hba-a1 mRNA after only 1-hour post-activation (approximately 2-fold), with an over 40-fold reduction after 24 hours (**Figure 3A**). These results were mirrored by activating T-lymphocytes with PMA/ionomycin (**Supplemental Figure 2A**), suggesting the downregulation of Hba-a1 is independent of T-lymphocyte receptor crosslinking. Next, we polarized CD4+ T- lymphocytes to T_H_1, T_H_2, T_H_17, and T_reg_ subtypes, and interestingly, all subtypes revealed a similar low Hba-a1 expression pattern, except for T_reg_ cells, which in fact upregulated Hba-a1 mRNA expression compared to naïve T-lymphocytes (**Figure 3B**). While polarization to T_reg_ cells requires supplementation of both transforming growth factor beta (TGF-β) and interleukin 2 (IL-2), the treatment of T-lymphocytes with either factor independently did not elevate Hba-a1 (**Supplemental Figure 2B**), suggesting the combination of factors is essential for Hba-a1 induction in T-lymphocytes. Furthermore, CD4+ memory T-lymphocytes (T_mem_) from wild-type unchallenged mice as well as CD8+ T-lymphocytes polarized to T_mem_ ex vivo also showed significant and robust elevations in Hba-a1 mRNA expression compared to their respective control T-lymphocytes (**Figure 3B- C).** Collectively, these data show that Hba-a1 expression significantly varies dependent upon polarization state of T-lymphocytes. One characteristic that may underly this complex pattern of Hba-a1 expression is metabolism, whereas T_H_0, T_H_1, T_H_2, T_H_17 cells (demonstrating low Hba-a1 levels) primarily rely on glycolysis for energy, while naïve, T_reg_, and T_mem_ cells (demonstrating high Hba-a1 levels) primarily rely on oxidative phosphorylation [29, 30, 44, 45]. To test how Hba-a1 may affect mitochondrial metabolism, Hba-a1 over- expression was induced via lentiviral transfection containing Hba-a1 under a constitutively active promoter in TK1 cells (**Supplemental Figure 2C**). Mitochondrial bioenergetics were assessed by the Seahorse mitochondrial stress test, and revealed that Hba-a1 over-expression led to an increase in oxygen consumption both at baseline and after FCCP injection, suggesting Hba-a1 increased mitochondrial metabolic activity (**Figure 3D**). Additionally, mitochondrial membrane potential, assessed by TMRE MFI, was significantly increased in cells over-expressing Hba-a1 compared to GFP controls, further supporting Hba-a1 acting in concert with mitochondrial metabolic activity (**Figure 3E**).

**Figure 3.**
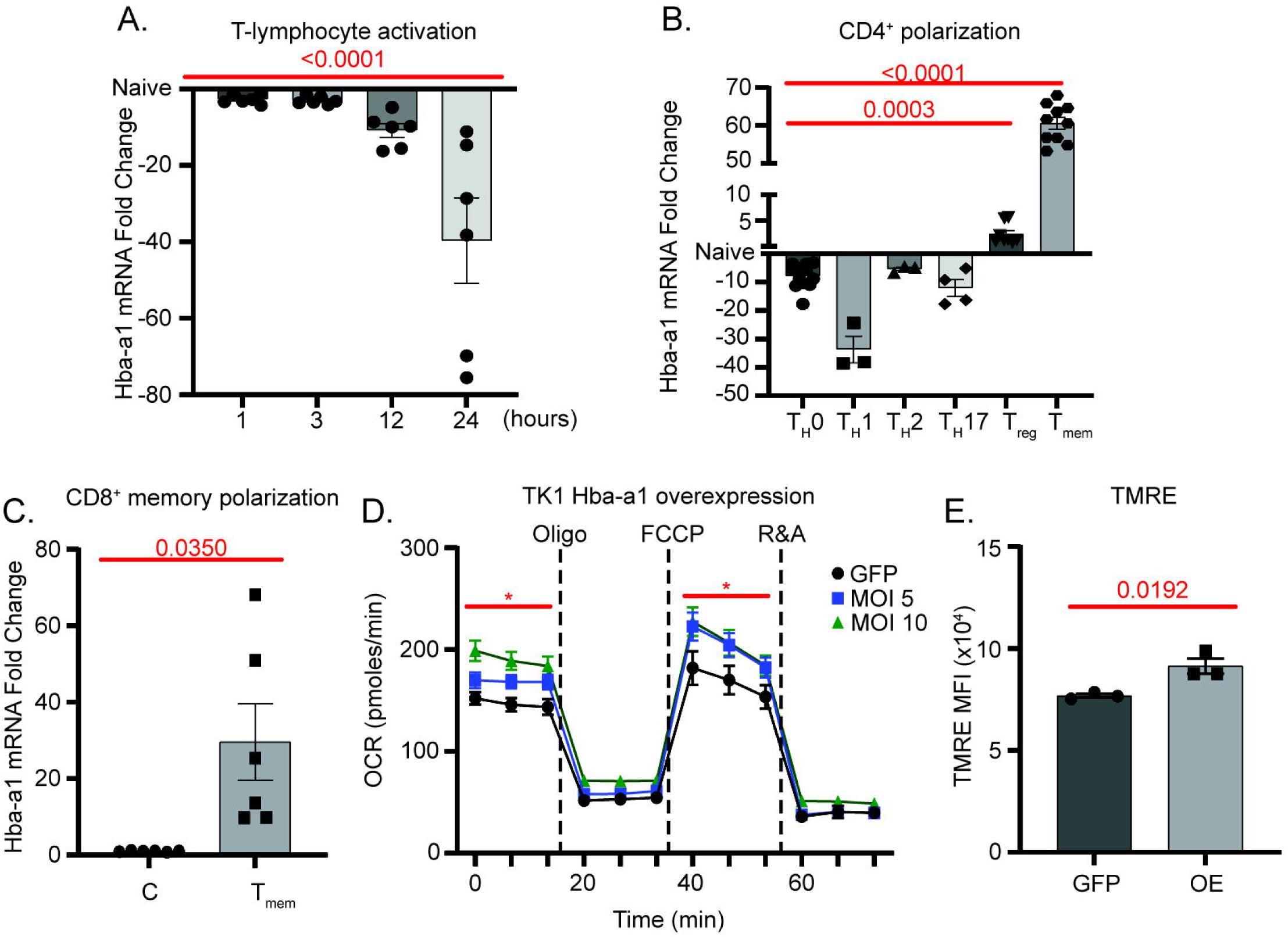
T-lymphocyte activation and polarization differentially alters Hba-a1 expression. **A.** Mouse naïve splenic T-lymphocytes activated with anti-CD3/28 Dynabeads at a 1:1 ratio and Hba-a1 mRNA assessed at time points indicated by RTqPCR. **B**. Hba-a1 mRNA assessment by RTqPCR of mouse naïve splenic CD4+ T-lymphocytes activated and polarized for 72 hours (as outlined in methods). **C**. Mouse naïve splenic CD8^+^ T- lymphocytes activated and polarized to memory T-lymphocytes (as outlined in methods) assessed for Hba-a1 via RTqPCR (C: IL-2 control, T_mem_: memory). **D**. TK1 cells transfected with GFP control lentivirus (GFP) or Hba-a1 lentivirus for 24 hours at various multiplicity of infections (MOI). Mitochondrial bioenergetics assessed by Seahorse mitochondrial stress test. **E**. Mitochondrial membrane potential of TK1 cells transfected with Hba- a1 lentivirus (MOI 5) or GFP control virus assessed by quantifying mean fluorescent intensity (MFI) of TMRE via flow cytometric analysis. Statistics by 1-way ANOVA (**A-B**), Student’s t-test (**C, E**), or 2-way ANOVA (**D**) with Tukey’s post-hoc analysis.

### Hba-a1 is transcriptionally regulated in naïve T-lymphocytes but silenced in glycolytic T-lymphocyte subtypes

To further explore the mechanism by which Hba-a1 is regulated within T-lymphocytes, TK1 cells were treated with H_2_O_2_ in the presence/absence of transcription inhibitors (i.e., actinomycin D or DRB) and Hba-a1 mRNA was assessed. Interestingly, both transcription inhibitors prevented the characteristic upregulation of Hba-a1 after H_2_O_2_ supplementation (**Figure 4A**), indicating that the redox-sensitivity of Hba-a1 appears to be transcriptionally regulated. Two well-known redox sensitive transcription factors, activator protein 1 (AP1) and nuclear erythroid 2 related factor 2 (NRF2) were examined as possible transcription factors responsible for the upregulation of Hba-a1 mRNA in response to redox modulators, but both yielded inauspicious results (**Supplemental Figure 3A-B**). Therefore, the redox-sensitive transcription factor responsible for Hba-a1 regulation in T-lymphocytes remains elusive. Curiously, while naïve T-lymphocytes demonstrated a prototypical upregulation of Hba-a1 in response to H_2_O_2_, activated T-lymphocytes had completely lost this redox-sensitivity (**Figure 4B**). This loss of responsiveness to redox agents as well as the rapid decrease in Hba-a1 mRNA levels after activation suggests the potential for higher level regulation, possibly at the chromatin level, in activated T-lymphocytes. To test this, chromatin accessibility of the putative Hba-a1 promoter region (∼1kb upstream of the transcription start site) was assessed in naïve and activated T-lymphocytes, and was found to be significantly more “closed” in activated T-lymphocytes (**Figure 4C**). The inaccessibility of the Hba-a1 locus in activated T-lymphocytes may underly the significant decrease of Hba-a1 in these cells as well as the loss of redox-sensitivity due to the inability of transcription factor binding. Overall, these data demonstrate a complex and dynamic regulation of Hba-a1 in T-lymphocytes that once again is highly dependent upon activation state of the cells.

**Figure 4.**
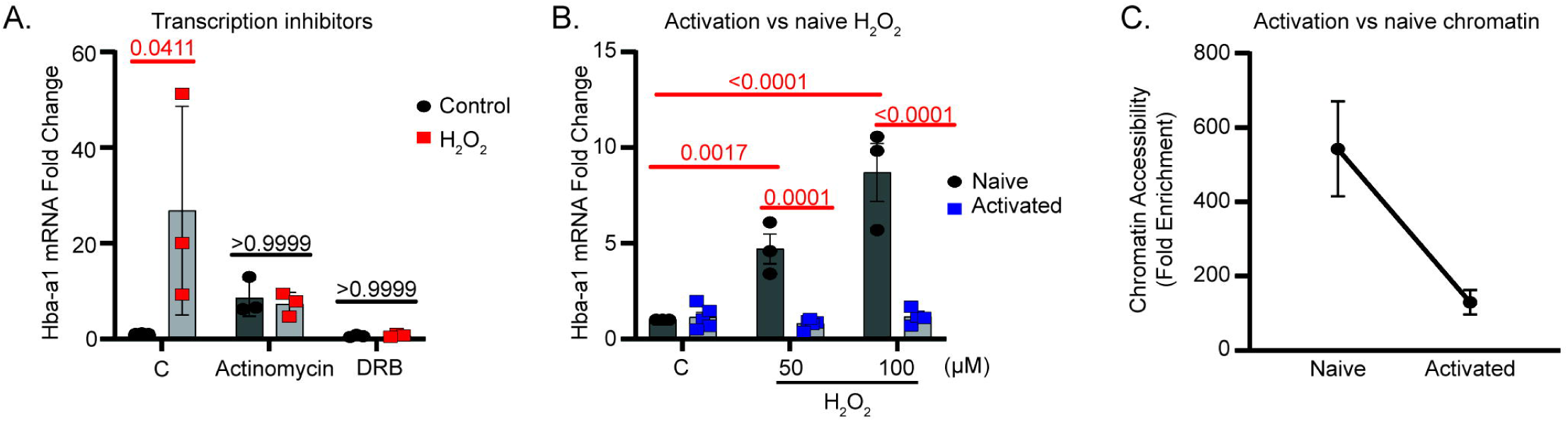
Hba-a1 is transcriptionally regulated in naïve T-lymphocytes but silenced in glycolytic T- lymphocyte subtypes. **A**. TK1 cells treated with actinomycin D (1µg/mL), or 5,6-dichloro-1-beta-D- ribofuranosylbenzimidazole (DRB, 100uM) for 30 minutes at 37**°**C, then treated ±250µM H_2_O_2_ for 6 hours. Hba- a1 mRNA assessed by RTqPCR (C: control). **B**. Mouse naïve and activated (with anti-CD3/28 beads for 48 hours) splenic T-lymphocytes treated ± H_2_O_2_ assessed for Hba-a1 mRNA by RTqPCR. **C**. Mouse naïve and activated splenic T-lymphocyte chromatin assessed for accessibility at the Hba-a1 promoter region (∼1000bp upstream of *Hba-a1* gene) by RTqPCR (n=5). Statistics by 2-way ANOVA with Tukey’s post-hoc analysis (**A**) or with Šídák’s post-hoc analysis (**B**).

## Discussion

While hemoglobin was once thought to be exclusively expressed in red blood cells, data from our lab and others demonstrate this ancient adage erroneous. Herein, our novel data supports the expression of hemoglobin alpha in T-lymphocytes, which transcriptionally responds to redox perturbations within the naïve form of these adaptive immune cells. These data further suggest that hemoglobin alpha functions as a mitochondrial modulating factor, given that the loss of this protein in T-lymphocytes led to a subsequent increase in mitochondrial ROS production and that Hba-a1 over-expression elevated mitochondrial respiration and membrane potential. Additionally, we show that Hba-a1 expression is greatly altered depending on T-lymphocyte differentiation, highlighting the intricate regulation of this protein. Together, these findings appear to only scratch the surface as to the complexity of hemoglobin function and regulation in T-lymphocytes, implying a completely unidentified mechanism that is essential in adaptive immune function.

Hemoglobin (both alpha and beta subunits) has now been identified in >10 cell types, however, the quest to define its function in these non-canonical cells has been a less successful venture. In our attempts to elucidate the function in T-lymphocytes, we, like other researchers have observed Hba-a1 upregulation in response to redox perturbations such as H_2_O_2_ [4, 15, 46], hypoxia [12, 14, 47], or auranofin. Interestingly, administration of NAC in the drinking water of RSDS mice prevented Hba-a1 induction in T-lymphocytes. A common theme among all of these redox agents is H_2_O_2_, suggesting this specific ROS may be responsible for the regulation of T-lymphocyte hemoglobin. However, the connection of Hba-a1 with mitochondria appears to be unique to T-lymphocytes, as opposed to many other cell types. For example, we observed Hba-a1 is upregulated in MnSOD knock-out mice, which produce significantly higher levels of mitochondrial ROS (both superoxide and H_2_O_2_) compared to WT mice [26]. Furthermore, mitochondrial-targeted catalase mice conversely show the opposite; the well-established increase in MitoSOX oxidation is lost along with the upregulation of Hba-a1 after RSDS. Since mitochondrial-targeted catalase specifically eliminates H_2_O_2_ from the mitochondria, this data further supports the role of this particular ROS in the regulation of Hba-a1 (while also reinforcing that H_2_O_2_ can oxidize fluorescent probes like MitoSOX). On the contrary, the loss of Hba-a1 in T-lymphocytes significantly potentiated the oxidation of MitoSOX after RSDS, suggesting Hba-a1 may function as a mitochondrial antioxidant. Together, these data propose a positive feedback loop between Hba-a1 expression and mitochondrial H_2_O_2_ levels. To date, subcellular localization of HBA in T-lymphocytes has proven challenging given the currently available antibodies, but identifying the specific location of HBA within T-lymphocytes may further illustrate this mitochondrial-associated role and remains an ongoing pursuit.

The balance of redox signaling in T-lymphocytes is crucial for proper cell activation and differentiation. Healthy, naïve T-lymphocytes have low levels of ROS, but shortly after T-cell receptor (TCR) stimulation, there is a controlled surge of ROS production to activate the nuclear factor of activated T cells (NFAT) transcription factor, and subsequently produce IL-2, which is essential for T-lymphocyte proliferation [25, 48]. Our group and others have demonstrated that mitochondrial ROS also contribute to this process, and that if the levels of these mitochondrial species are too low or too high, T-lymphocyte activation and function is compromised [24, 25].

Differentiation into various effector cells is also mediated in part by ROS levels which may have dire consequences if not carefully controlled, such as an increased risk for autoimmune disorders [48]. Thus, it is imperative that T-lymphocytes have multiple layers of redox control to maintain the desired activation and polarization state. Unlike the well-studied antioxidant glutathione (GSH) in T-lymphocytes, which is upregulated after T-lymphocyte activation to buffer the increase in ROS production [49, 50], we demonstrate that Hba-a1 mRNA is rapidly decreased after only 1 hour post-stimulation, and is essentially undetectable after 24 hours.

This stark decrease suggests Hba-a1 must be silenced to properly activate and enter a glycolytic metabolic state, as opposed to the oxidative phosphorylation/fatty acid oxidation metabolic environment of naïve, T_reg_, and memory T-lymphocytes where Hba-a1 is significantly upregulated [51]. We observed this decrease of Hba- a1 in glycolytic T-lymphocyte subtypes may be due to rapid epigenetic remodeling that occurred in the region directly upstream of Hba-a1 (∼1000bp) within 24 hours post activation. Moreover, the lack of redox sensitivity of Hba-a1 induction in activated T_H_0 cells 48 hours post-activation further suggests chromatin inaccessibility.

This quick condensation and decrease in Hba-a1 expression postulates that Hba-a1 may also function as a critical mediator of T-lymphocyte activation that is possibly dependent upon metabolic state.

As previously mentioned, hemoglobin’s well-known function as an oxygen carrier has been well established for roughly a century, thus there is a breadth of data examining hemoglobin regulation in red blood cells.

Unfortunately, known inducers of hemoglobin such as erythropoietin and hemin did not induce Hba-a1 expression in T-lymphocytes, suggesting a differential regulatory mechanism compared to its canonical predecessor. This is not all too surprising, given that mature red blood cells lack a nucleus. Even transforming growth factor beta (TGFβ), sometimes known as “a master regulator of T-lymphocytes” did not induce Hba-a1 upregulation on its own in T-lymphocytes, despite observations of HBA expression after TFGβ supplementation in K562 leukemia cell line [52, 53]. Similarly, there are several known transcription factors that upregulate Hba- a1 in developing red blood cells, mainly GATA1 and KLF2/4 [54, 55]. In a couple of studies, these factors did appear involved in Hba-a1 regulation in non-canonical cell types [11, 56], but it remains unclear if these play a role in Hba-a1 regulation in T-lymphocytes. Other researchers suggest that hypoxia inducible factor alpha (HIF1α) directly acts as the transcription factor for Hba-a1 [6, 12]. This is unlikely to be the sole transcription factor of Hba-a1 in T-lymphocytes, because while hypoxia did induce Hba-a1 expression, many other redox perturbations and polarizations states also induced Hba-a1 in the absence of upregulation of canonical HIF1α response genes (i.e. NAD(P)H Quinone Dehydrogenase 1 (Nqo1), data not shown). Given that other redox-sensitive transcription factors, Nrf2 and AP1, also failed to alter Hba-a1 expression, at this time the master transcriptional regulator of T-lymphocyte Hba-a1 remains elusive.

Interestingly, throughout evolution, gene duplication of hemoglobin alpha led to the creation of two identical hemoglobin alpha copies (*Hba-a1* and *Hba-a2* in mice; *HBA1* and *HBA2* in humans), which have evolutionarily co-existed cis-chromosomally ever since. These two copies differ primarily in the untranslated regions of the gene, and translate into 100% identical protein sequences. This evolutionary ‘built-in backup’ of hemoglobin alpha highlights the importance of this protein; in fact, people missing two of the four hemoglobin alpha genes (4 total given duplicate chromosomes) will likely have mild to no symptoms due to the compensation from two healthy copies of the gene [57]. This compensation, however, suggests that the phenotypes seen in our current Hba-a1 knock-out animals may be severely diminished, thus partially masking hemoglobin alpha’s potential indispensable role as a mitochondrial antioxidant in T-lymphocyte biology. Hence, our laboratory has created a double floxed Hba-a1/Hba-a2 knock-out animal to address this potential compensation, and will be examining how T-lymphocyte mitochondrial function, activation, and polarization is affected by the loss of both copies of hemoglobin alpha. Additionally, we have also detected hemoglobin beta subunits expressed in murine T-lymphocytes. Interestingly, the beta subunits do not appear to act like the alpha subunits, and are more consistently expressed across all states of T-lymphocyte activation and differentiation (as well as do not appear to be redox sensitive). It remains unknown if hemoglobin beta is expressed at the protein level in T-lymphocytes like the alpha subunits, or if the two proteins even interact. This observation only further confirms the complexity of hemoglobin expression T-lymphocytes that remains to be uncovered.

In addition to the conserved duplication of hemoglobin alpha, the genes surrounding hemoglobin alpha locus have also been on the same genetically-conserved timeline. The hemoglobin alpha cluster has existed downstream of the *Nprl3* gene dating back to the jawed vertebrates. Interestingly, it has been shown that the introns within this gene act as upstream enhancers for hemoglobin alpha [58–61]. Furthermore, the *Nprl3* gene itself encodes a subunit in the protein complex GATOR1, which negatively regulates mTOR signaling. mTOR, or mammalian target of rapamycin, is a protein kinase that is involved in mitochondrial oxygen consumption [62] as well as T-lymphocyte activation, polarization, and memory formation [63]. In naïve T-lymphocytes, mTOR is basally active, whereas activation leads to robust mTOR stimulation [64]. Interestingly, T-lymphocytes that lack mTOR cannot polarize to glycolytic-dependent T_H_1, T_H_2, or T_H_17 subtypes, and subsequently will only polarize to T_reg_ cells [65]. The level of mTOR signaling appears in direct alignment with the expression pattern of Hba-a1 we demonstrate in activated T-lymphocytes. Taken in combination with the preservation of this chromosomal arrangement for millions of years, the tight correlation between mTOR, Hba-a1 expression, and T-lymphocyte bioenergetics during polarization warrants further investigation of this potential regulatory mechanism.

Last, about 7% of the world’s population are carriers for hemoglobinopathies, creating an enormous and global disease burden [66]. This class of genetic diseases is classified by a mutation in one or more hemoglobin subunits and includes sickle cell anemia, alpha, and beta thalassemia. Interestingly, there is a strong correlation of patients with hemoglobinopathies developing autoimmune disorders such as rheumatoid arthritis and psoriatic arthritis, as well as systemic lupus erythematosus [67–69]. While it is not completely known what is leading to this increased risk of autoimmunity in hemoglobinopathy patients, the potential role of mutated hemoglobin in adaptive immune cells should be investigated as one potential causal link between these two maladies. Our data herein suggest that the loss of hemoglobin function may lead to a more activated and pro-inflammatory state in T-lymphocytes, which may underly these unforeseen physiological consequences of hemoglobinopathies and provide new therapeutic strategies for these patients or others with adaptive immune system disorders.

## Acknowledgements

### Funding sources

This work was supported by the National Institutes of Health (NIH) R01HL158521 (AJC). We thank Drs. Holly Van Remmen and Brant Isakson for their contributions of mouse models used in this work. We thank Texas A&M University’s Flow Cytometry and Cell Sorting Facility for the use of their seahorse bioanalyzer, as well as Texas A&M University’s Preclinical Phenotyping Core for the use of the Protein Simple Jess Automated Western Blot.

## Author contributions

ECR and AJC designed research studies. All authors conducted experiments, acquired data, and/or performed analyses. ECR and AJC wrote the manuscript, while all authors approved the final version of the manuscript. AJC provided funding and experimental oversight.

## Abbreviations

Hba-a1, hemoglobin alpha-a1; MnSOD KO, manganese superoxide dismutase knock-out; WT, wild-type; RSDS, repeated social defeat stress; LPS, lipopolysaccharide; PCR, polymerase chain reaction; ROS, reactive oxygen species; RTqPCR, real time quantitative polymerase chain reaction; H_2_O_2_, hydrogen peroxide; superoxide, O_2_^•-^ ; AFN, auranofin; DRB, 5,6-Dichlorobenzimidazole 1-β-D-ribofuranoside; MFI, mean fluorescence intensity; mTOR, mammalian target of rapamycin; PMA, phorbol myristate acetate

