## Supplemental Data for "Hemoglobin alpha is a redox-sensitive mitochondrial-related protein in T-lymphocytes"

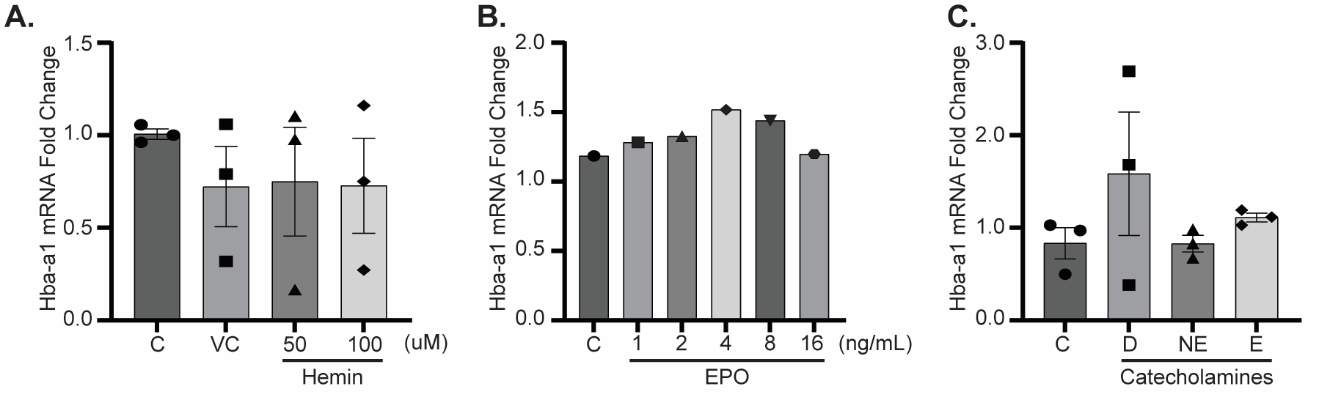

**Supplemental Figure 1. Treatments that do not induce Hba-a1 in T-lymphocytes. A**. TK1 cells treated with vehicle control (VC) or hemin for 24 hours, then assessed for Hba-a1 mRNA expression by RTqPCR (C: control). **B**. TK1 cells treated with erythropoietin (EPO) for 24 hours assessed for Hba-a1 mRNA expression by RTqPCR (C: control). **C**. TK1 cells treated with 10µM of dopamine (D), norepinephrine (NE), or epinephrine (E) for 24 hours assessed for Hba-a1 expression via RTqPCR. (C: control). Statistics (not significant, not shown) by 1-way ANOVA (**A-C**).

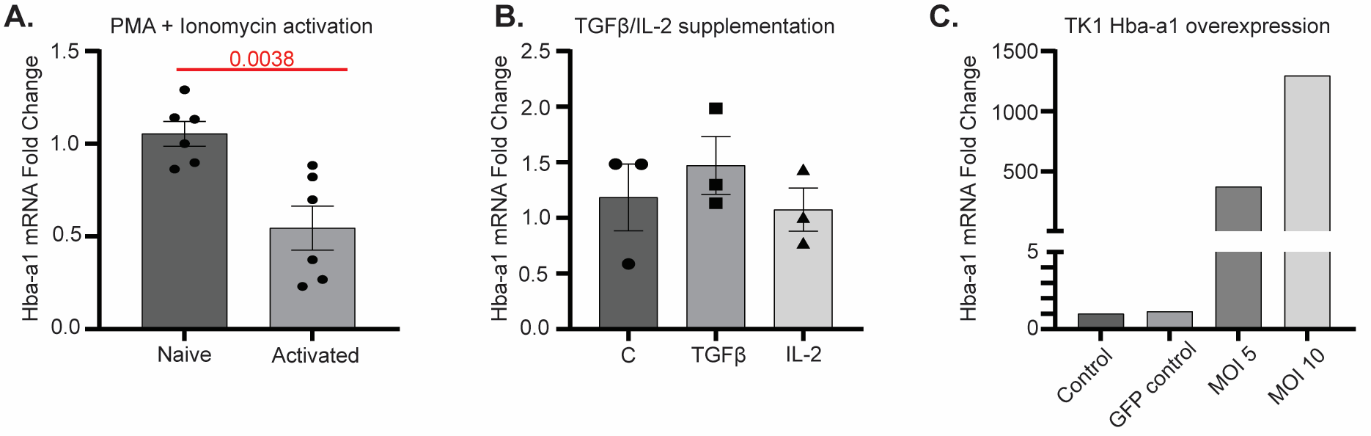

**Supplemental Figure 2. A.** Mouse naïve splenic T-lymphocytes activated in the presence of phorbol myristate acetate (PMA) and ionomycin for 24 hours and Hba-a1 mRNA expression assessed by RTqPCR. **B**. TK1 cells treated with 15 ng/mL transforming growth factor beta (TGFβ) or 150 ng/µL IL-2 assessed for Hba-a1 expression via RTqPCR. (C: control). **C**. TK1 cells transfected with GFP control virus (GFP control) or Hba-a1 overexpression virus (MOI 5, MOI 10) assessed for Hba-a1 mRNA by RTqPCR. Statistics by Student’s t-test (**A**) or 1-way ANOVA (**B-C**, not significant, statistics not shown).

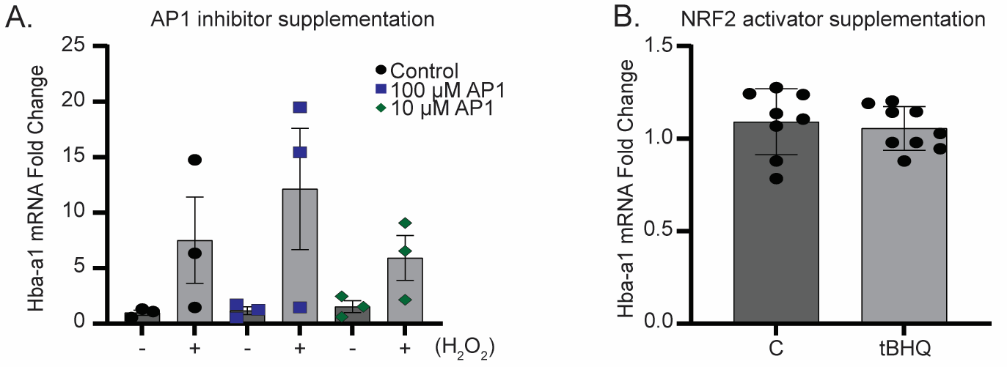

**Supplemental Figure 3. Known redox transcription factors that do not appear to increase Hba-a1 mRNA expression in T-lymphocytes. A**. TK1 cells treated with activator protein-1 (AP1) inhibitor T-5224 incubated for 30 minutes and treated with ± 250µM H_2_O_2_ for 6 hours. Hba-a1 mRNA assessed via RTqPCR. **B**. TK1 cells treated with 5 µM tert-butylhydroquinone (t-BHQ) for 6 hours at 37°C assessed for Hba-a1 mRNA by RTqPCR. Statistics (not significant, not shown) by 2-way ANOVA (**A**) or Student’s t-test (**B**).
